# Neural network modeling of differential binding between wild-type and mutant CTCF reveals putative binding preferences for zinc fingers 1-2

**DOI:** 10.1101/2021.09.23.461552

**Authors:** Irene M. Kaplow, Abhimanyu Banerjee, Chuan Sheng Foo

## Abstract

**Background:** Many transcription factors (TFs), such as multi zinc-finger (ZF) TFs, have multiple DNA binding domains (DBDs) with multiple components, and deciphering the DNA binding motifs of individual components is a major challenge. One example of such a TF is *CCCTC-binding factor* (CTCF), a TF with eleven ZFs that plays a variety of roles in transcriptional regulation, most notably anchoring DNA loops. Previous studies found that CTCF zinc fingers (ZFs) 3-7 bind CTCF’s core motif and ZFs 9-11 bind a specific upstream motif, but the motifs of ZFs 1-2 have yet to be identified.

**Results:** We developed a new approach to identifying the binding motifs of individual DBDs of a TF through analyzing chromatin immunoprecipitation sequencing (ChIP-seq) experiments in which a single DBD is mutated: we train a deep convolutional neural network to predict whether wild-type TF binding sites are preserved in the mutant TF dataset and interpret the model. We applied this approach to mouse CTCF ChIP-seq data and, in addition to identifying the known binding preferences of CTCF ZFs 3-11, we identified a GAG binding motif for ZF1 and a weak ATT binding motif for ZF2. We analyzed other CTCF datasets to provide additional evidence that ZFs 1-2 interact with the motifs we identified, and we found that the presence of the motif for ZF1 is associated with Ctcf peak strength.

**Conclusions:** Our approach can be applied to any TF for which *in vivo* binding data from both the wild-type and mutated versions of the TF are available, and our findings provide an unprecedently comprehensive understanding of the binding preferences of CTCF’s DBDs.

## BACKGROUND

Mutations of individual DNA binding domains (DBDs) within transcription factors (TFs) have been associated with developmental disorders [1, 2] and bleeding disorders [3], and differences between species in individual DBDs within TFs have been associated with species-specific gene expression [4] and speciation [5]. Yet, even though DNA binding motifs of thousands of metazoan TFs have been characterized, many TFs have multiple DNA binding domains (DBDs) whose specific binding preferences are unknown. In fact, the most common TF family in humans, Cys2His2 (C2H2) zinc finger (ZF) TFs [6, 7], consists of TFs with multiple ZF DBDs, and many of these ZFs’ individual binding preferences have not been investigated.

A previous study investigated the binding preferences of ZFs within C2H2 ZF TFs by doing *in vitro* Bacterial 1-Hybrid (B1H) assays of over 160,000 ZFs [8] to determine the individual 3bp [9] binding preferences of each ZF. The study then presented a machine learning model trained on this data to predict the position weight matrices (PWMs) of C2H2 ZF TFs. Unfortunately, for the majority of TFs, less than two thirds of PWM columns were predicted correctly, demonstrating the limitations of using *in vitro* assays of individual DBDs to determine binding preferences of DBDs within a full TF. Another study described how DBDs can influence each other’s binding within the context of a TF [10], further illustrating the limitations of studying binding preferences of individual DBDs out of context.

To identify the binding preferences of DBDs within a TF within the context of the other DBDs, previous studies have introduced loss-of-function mutations within specific DBDs, assayed the sequences to which the mutants bind, and used the results of the assay to determine the specific components of TFs’ motifs that interact with a DBD [1, 11, 12]. In particular, one of these studies induced loss-of-function histidine-to-arginine mutations separately each of the 11 ZFs of mouse *CCCTC-binding factor* (CTCF), a C2H2 ZF TF that has been implicated in diverse roles in transcriptional regulation [13, 14] due to its ability to anchor DNA loops [15, 16], and did ChIP-seq on each mutant [11]. The study found that ZFs 3 through 7 interact with part of CTCF’s known core motif, a 19 base-pair sequence that has been shown to bind CTCF in many studies [17], and that ZFs 8 through 11 interact with an upstream motif that had been identified by a few earlier studies [18–20] (**Supplemental Figure 1**), demonstrating the viability of assaying binding of mutated TFs to understand individual DBD binding preferences. These findings were supported by additional studies; one study used CTCF deletions to show that only ZFs 4 through 7 interact with base pairs 4 through 15 of its core motif [21], and another study used electrophoretic mobility shift assays (EMSA) of CTCF with parts of its motif mutated to suggest that ZF 7 or 8 binds to base pairs four through six of the core motif [22]. However, despite this vast body of work, the *in vivo* binding preferences of ZFs 1 and 2 remain unknown. In addition, a recent study showed that mutations in ZFs 1 and 10 disrupt DNA loops [23], and another recent study showed that CTCF-s, a CTCF isoform that does not have ZFs 1-3, is unable to interact with cohesin [24], underscoring the potential value in understanding the ways that CTCF’s ZFs that do not bind to the core motif interact with DNA.

To better leverage *in vivo* experiments of mutated TFs to decipher the binding preferences of individual DBDs, we developed a novel approach to analyzing the data from mutant TF ChIP-seq experiments [11] in which, in contrast to the earlier study, which did *de novo* motif discovery on the sequences within the peaks from wild-type CTCF and then scanned the peaks from the mutated CTCF for the discovered motifs [11], we directly leverage the differences between the wild-type and mutant datasets. We do this by setting up a differential peak prediction task, in which we train a deep convolutional neural network [25, 26] to use DNA sequence to predict whether a peak from wild-type TF ChIP-seq is preserved in the mutant dataset or is significantly stronger in the wild-type dataset. Our intuition is that, if a model can predict whether a peak is significantly stronger in the wild-type dataset than in the mutant dataset, then the model should have learned sequence patterns related to the binding preferences of the mutated DBD, and interpreting the model should reveal these binding preferences.

We applied our approach to the CTCF mutant ChIP-seq datasets [11] and interpreted what each model learned to identify motifs associated with each ZF. The interpretations recapitulated earlier findings about which ZFs interact with the core and upstream motifs and identified a novel downstream motif, GAGCCA, that may be bound by ZF 1, as well as a putative weak interaction between ZF 2 and the sequence ATT. We found that the core motif followed by our discovered downstream motif occurs in CTCF HT-SELEX reads from the final cycle, that the core motif followed by the discovered occurs more frequently in CTCF ChIP-seq peaks that do not overlap CTCF-s ChIP-seq peaks than in those that do overlap CTCF-s ChIP-seq peaks, and that the discovered downstream motif matches *in vitro* data-based computational predictions of the ZF 1 motif and has been shown to bind CTCF in a previous EMSA study. We also found that the presence of the discovered downstream motif is correlated with CTCF peak strength. Our approach can be applied to any TF with multiple DBDs for which wild-type and mutated DBD *in vivo* binding data are available, and our results from applying our approach to CTCF provide the first insights into the *in vivo* binding preferences of CTCF’s most downstream ZFs.

## RESULTS

### Putative Motifs of CTCF’s Zinc Fingers Identified by Interpreting Wild-Type versus Mutant Differential Peak Prediction Models

To identify motifs related to the binding of each ZF in CTCF, we trained and interpreted a neural network for predicting whether a peak would be significantly weaker according to DESeq2 [27] in the mutant dataset than in the wild-type dataset (**Methods**, **Supporting Website**). We trained a separate model for each ZF mutant ChIP-seq dataset from [11] (**Supplemental Figure 1**) and, upon finding that our models had good performance, used deepLIFT with the Rescale rule [28] followed by TF-MoDISco [29] to identify motifs (called “TF-MoDISco motifs”) that the model had learned (**Figure 1**, **Methods**, **Supporting Website**). We identified putative motifs for all of the ZFs in CTCF.

**Figure 1:**
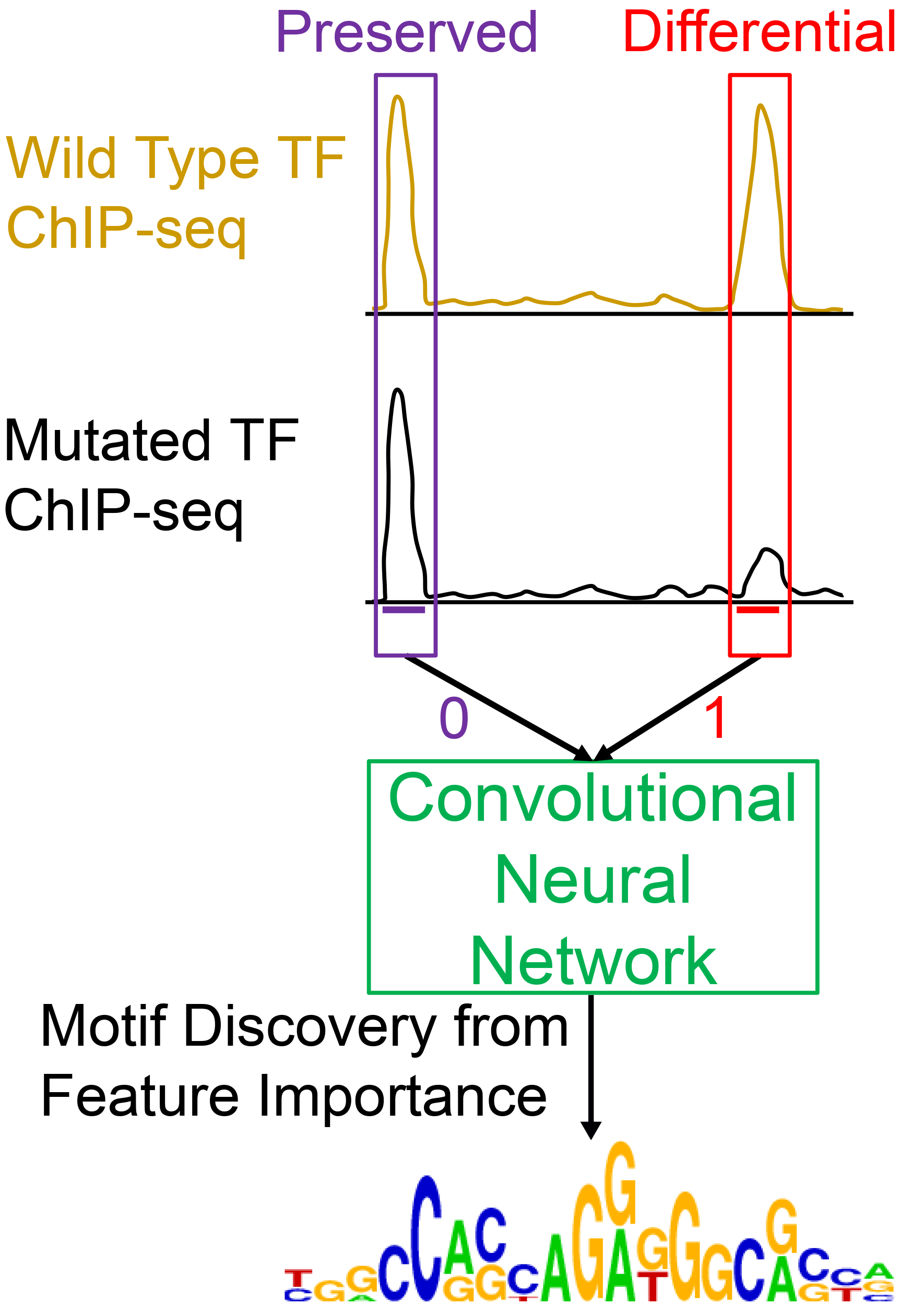
Using Differential Peak Prediction to Identify Motifs of Different DNA Binding Domains. To identify the motif of a DBD, we train a deep convolutional neural network to predict whether a TF ChIP-seq peak is preserved or significantly weaker in a dataset from that TF with a mutated DBD relative to a dataset from the wild-type TF. We then use deepLIFT followed by TF-MoDISco to identify the motifs that the neural network learned. The motif logo in this figure is the Hocomoco human CTCF motif downloaded from CIS-BP [17].

### Neural Network Outperforms Models with Original Motif Hit Scores as Features

To evaluate our neural network and the TF-MoDISco motifs, we compared three approaches for predicting whether a CTCF peak would be substantially weaker in a mutant CTCF dataset: our neural networks, logistic regressions with the motif hit score of the best TF-MoDISco motif hit as the feature, and logistic regressions with motif hit scores of motif hits from [11] as features. We found that all models performed well for ZFs 3-7, but our neural networks and the logistic regressions with the TF-MoDISco motif hit score alone had substantially better performance than the logistic regressions with the original motif hit scores for the other ZFs (**Figure 2a**). For ZFs 8-11, we also compared the performances of our neural networks and the logistic regressions with the TF-MoDISco motif hit score to the performances of the score from single motif consisting of the original upstream motif followed by five base pairs (the most common spacing found in [11]) followed by the original core motif; these logistic regressions’ performance were comparable to those of the logistic regressions with the TF-MoDISco motif hit score for ZFs 9-11 and worse than other methods for ZF 8 (**Supplemental Figure 2**).

**Figure 2:**
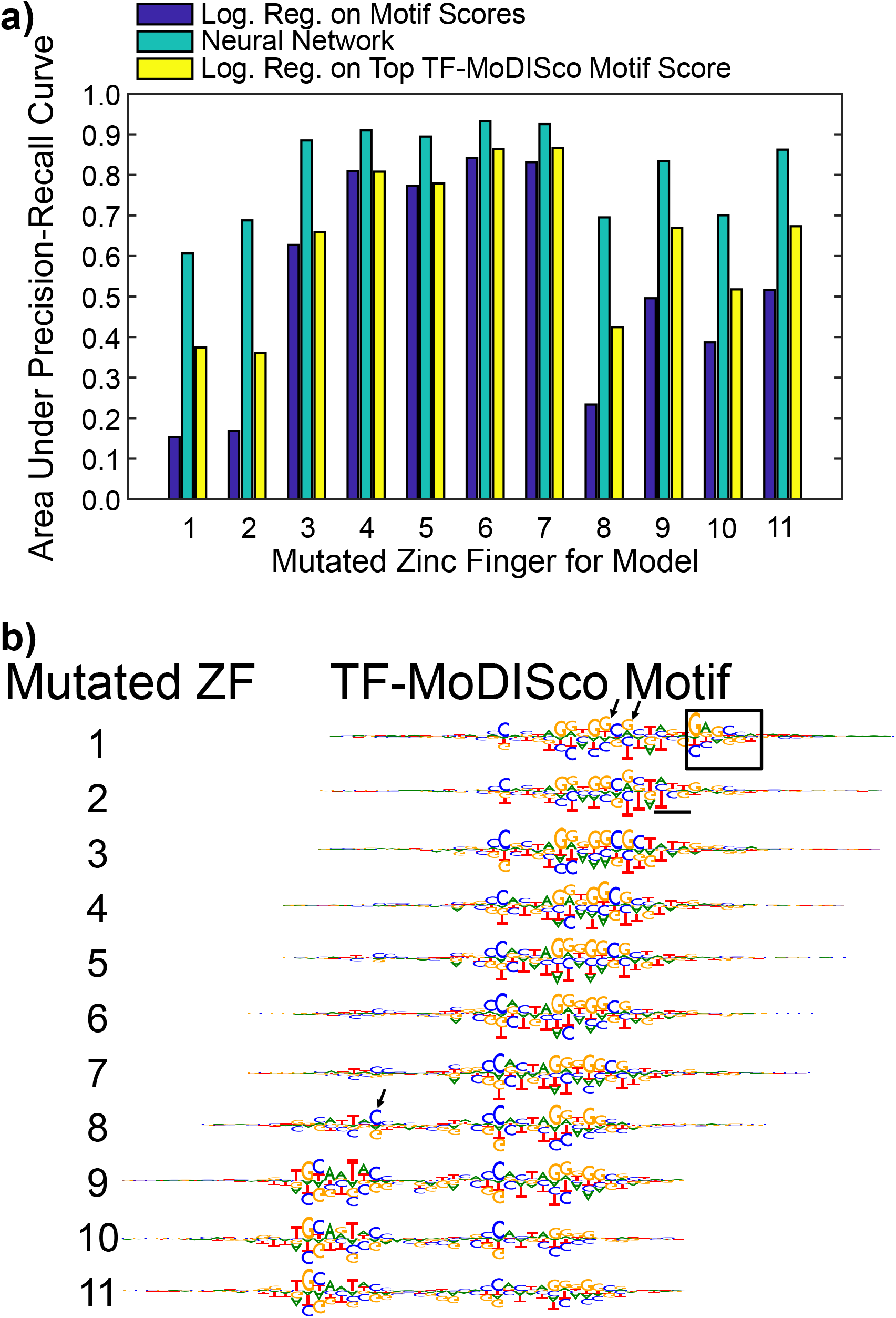
Performance of Neural Networks. **a)** We compared the performance of our neural networks to those of logistic regressions in which the features were the motif hit scores of the motifs from [11]. We also compared both of these models to logistic regressions with the top TF-MoDISco motif hit scores as their only features. Performance was measured by the area under the precision-recall curve (AUPRC). **b)** We aggregated the hypothetical scores of the seqlets corresponding to the motifs from deepLIFT followed by TF-MoDISco to visualize the TF-MoDISco motifs. The box indicates the discovered downstream motif, and the underlined part indicates the weak putative motif for ZF 2. The TF-MoDISco motif for ZF 1 has a G or a T at a position where the other TF-MoDISco motifs have a G (indicated by first arrow) and a G or an A at a position where the other TF-MoDISco motifs have a G (indicated by second arrow). The TF-MoDISco motif for ZF 8 emphasizes a downstream nucleotide in the upstream motif (indicated by arrow).

### Important Features Learned by Neural Networks Include Known Motifs for Zinc Fingers 3-11 and Novel Motifs for Zinc Fingers 1-2

#### Interpreting Neural Networks Revealed Known CTCF Motifs

We compared the TF-MoDISco motifs to known CTCF motifs. We found that our neural network learned motifs similar to the known core motif as being indicative of a stronger peak in the wild-type for ZFs 3-7 and motifs similar to the known upstream motif as being indicative of a stronger peak in the wild-type for ZFs 8-11, which is consistent with the findings of [11] (**Figure 2b**). Previous studies identified the five base pair spacing in our top TF-MoDISco motif as the most common spacing between the core and upstream motifs but also found that a six base pair spacing occurred frequently [11, 18–20, 30]. We therefore investigated all of the TF-MoDISco motifs for each neural network (**Supporting Website**) and, for ZFs 9-11, found that the second highest-ranked TF-MoDISco motif (the TF-MoDISco motif with the second highest number of supporting seqlets) was the upstream motif, followed by six base pairs, followed by the core motif (**Supplemental Figure 3**).

#### Interpreting Neural Networks Revealed Novel Motifs for ZFs 1-2 Confirmed by CTCF HT-SELEX Data

When identifying the important sequences for the neural networks for the mutants of ZFs 1-2, we discovered a novel downstream GAGCCA motif occurring 2bp downstream of the core motif and a weaker ATT motif connecting the core and discovered downstream motif as being indicative of a stronger peak in the wild-type (**Figure 2b**, **Supplemental File 1**). To investigate if CTCF can bind these motifs, we re-analyzed published HT-SELEX data for CTCF [31] to determine if there is an enrichment of reads containing the core followed by the discovered downstream motif in cycle 4 (final round) relative to cycle 0 (control) (**Methods**). First, to evaluate the reliability of this approach, we did this for the core motif only and found a significant enrichment (p = 0.0) (**Supplemental Figure 4**). We then found an enrichment for the core motif followed by the discovered downstream motif (p = 1.17 x 10^-245^) (**Figure 3b**). In fact, the HT-SELEX reads with the best matches to the core motif followed by the downstream motif (FIMO q-value < 0.001) have ATT connecting the two motifs (**Supplemental Figure 5**), which is the putative motif that we found for ZF 2.

**Figure 3:**
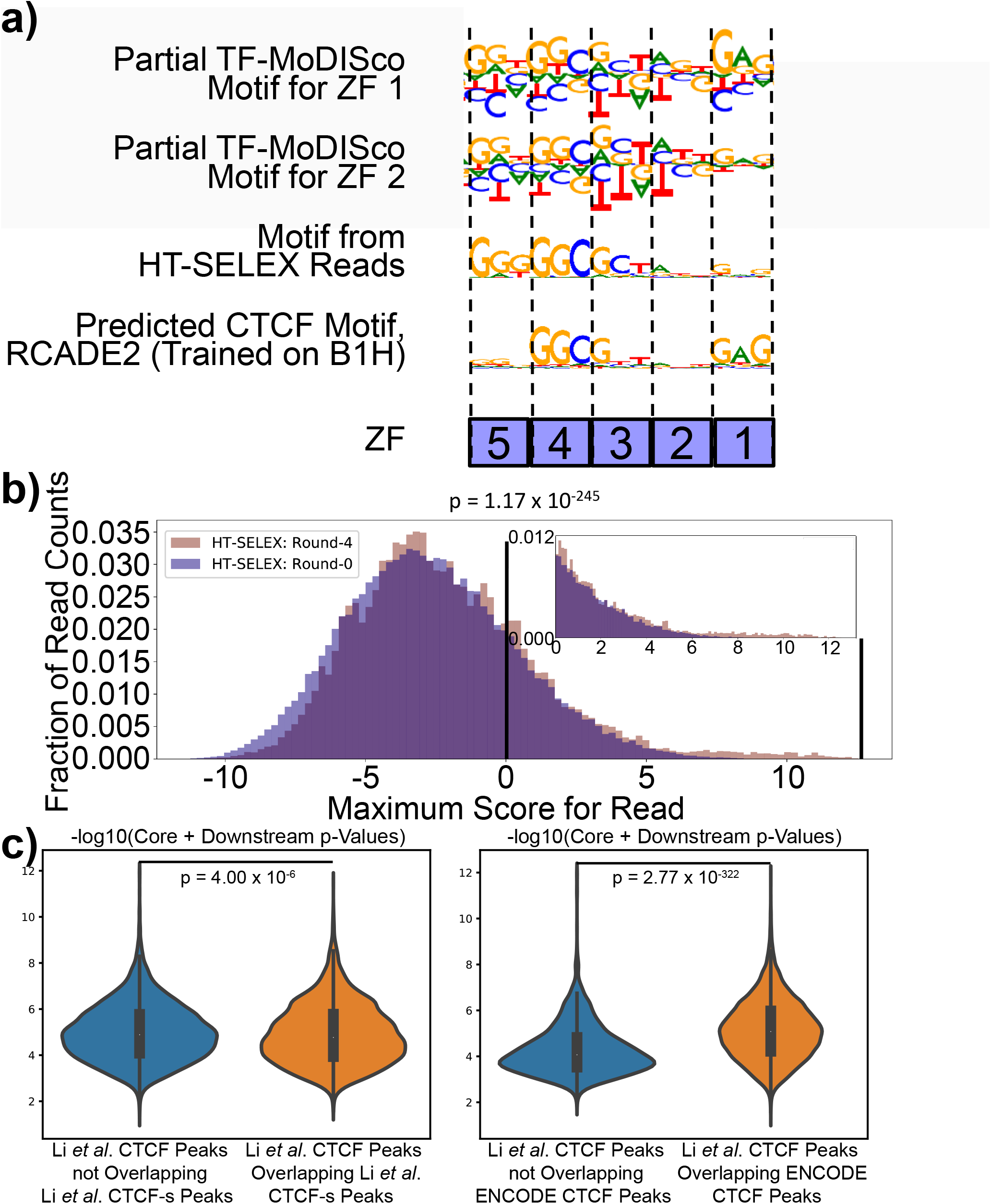
Comparisons of Discovered Downstream Motif to Other CTCF Data. **a)** We compared our TF-MoDISco motifs from the mutants of ZFs 1 and 2 to aggregated reads from CTCF HT-SELEX cycle 4 and to computationally predicted motifs of CTCF’s DBDs from the RCADE2 model, which was trained on *in vitro* B1H ZF binding data. **b)** We compared motif matches of the core followed by discovered downstream motif in reads from CTCF HT-SELEX data in cycle 0 to cycle 4. **c)** We compared the strength of the core followed by the downstream motif in HeLa cell *CTCF* peaks from [24] to HeLa peaks from CTCF’s alternative isoform from the same study and HeLa CTCF peaks from ENCODE [32].

#### Discovered Downstream Motif Is Associated with Lack of CTCF-s Binding

We also compared the p-values of the motif hits for the core followed by the discovered downstream motif in HeLa cell ChIP-seq peaks for CTCF and CTCF-s – the alternative isoform of CTCF that is missing ZF 1, ZF 2, and part of ZF 3 – from [24]. We found that these p-values were significantly lower (negative log base ten of the p-values was significantly higher) for the CTCF peaks that do not overlap CTCF-s peaks than they were for the CTCF peaks that do overlap CTCF-s peaks (p = 4.00 x 10^-6^), suggesting that the lack of ZFs 1-3 is associated with a lack of binding to the downstream motif. To investigate whether this result could be explained by the core motif followed by the downstream motif occurring more frequently in CTCF binding sites that are less reproducible across experiments, we downloaded HeLa cell CTCF ChIP-seq peaks from ENCODE [32] and compared the core followed by the discovered downstream motif hit p-values for the CTCF ChIP-seq peaks from [24] that overlap the ENCODE CTCF ChIP-seq peaks to those that do not. For this comparison, we found a significant trend in the opposite direction (p = 2.77 x 10^-322^) (**Figure 3c**). These results suggest that the lack of the core followed by the discovered downstream motif is associated with the lack of binding of CTCF ZFs 1-3.

#### Discovered Downstream Motif Has Supporting Evidence from Previous CTCF Studies

We obtained additional evidence that our discovered downstream motif interacts with CTCF. The most non-degenerate part of this motif (GAG) is almost identical to the computationally predicted motif for ZF 1 according to multiple models that were trained on *in vitro* ZF binding data from B1H assays (**Figure 3a, Supplemental Figure 6**), suggesting that ZF 1 interacts with this downstream motif [33–35]. In addition, a recent study showed that the upstream four nucleotides of this downstream motif are found at CTCF sites in the mouse IgH locus; this study did EMSA on multiple variants of the CTCF motif including two variants containing these upstream four nucleotides and found that CTCF was able to bind both variants [36]. Furthermore, the downstream 3bp of this motif (CCA) is similar to the upstream 3bp of the 4bp downstream motif identified in CTCF-cohesin co-binding sites in [37]. In spite of this evidence suggesting the existence of our downstream motif, this motif has not been previously shown to directly interact with CTCF ZF 1 *in vivo*.

#### *Neural Networks’ Nucleotide-Level Relative Importance Scores Reveal Putative Combinatorial Binding Preferences that Were Supported by* in vitro *TF Binding Assays*

Our neural networks’ differences between relative importance scores of nucleotides in motifs for different ZFs provided potential insights into additional differences between CTCF’s binding preferences when different ZFs interact with DNA. For example, the TF-MoDISco motif for the mutant of ZFs 1-2 had a degenerate position in the core motif that could be a G or a T and another that could be a G or an A. In contrast, the TF-MoDISco motifs for the mutants of ZFs 3-7 placed a substantially stronger importance on the G than the T in the first position, and the TF-MoDISco motifs for the mutants of ZFs 8-11 placed no importance on the T in that position. Interestingly, the EMSA done on the core followed by the discovered downstream motif had a T in that position [36], showing that CTCF can bind to the core motif when there is a T in that position and the discovered downstream motif is present. Likewise, the TF-MoDISco motifs for the mutants of ZFs 3-11 placed no importance on the G in the second position, providing another example of degeneracy being tolerated in the presence of only the downstream motif (**Figure 2b**).

The neural networks for ZF 8 also placed higher importance on the downstream nucleotides of the upstream motif than did the neural networks for other ZF mutants (**Figure 2b**). Additionally, the neural networks for ZFs that are thought to interact with upstream parts of the core motif placed a weak importance on the upstream motif, while the neural networks for ZFs that are thought to interact with downstream parts of the core motif placed a weak importance on the downstream motif. Thus, in addition to identifying putative *in vivo* motifs for ZFs 1-2, and our neural networks provided insights into the relative importance of various parts of motifs for interactions between DNA and different ZFs.

### Presence of Discovered Downstream Motif Is Associated with CTCF Peak Strength

Since the discovered downstream motif is present in only a strict subset of wild-type CTCF peaks, we investigated whether the presence of this motif is associated with other differences between CTCF peaks. We identified occurrences of the upstream, core, and discovered downstream motifs and combinations of these motifs in the wild-type CTCF peaks using FIMO [38] (**Supporting Website**). We then examined the relationship between motif presence and peak strength (**Methods**). We found that peaks with the core motif and the discovered downstream motif tend to be stronger than peaks with the core motif that do not have the discovered downstream motif (p = 3.24 x 10^-106^). Since the presence of a motif in a peak is dependent on the motif hit cutoff, we also applied a stricter threshold for the motif hit cutoff and obtained a similar result (p = 7.77 x 10^-89^). In addition, we evaluated the relationship between the motif hit q-values of the different motif combinations and the peak signal and found that the correlation is significantly stronger for the core motif followed by the discovered downstream motif than it is for the core motif alone (p = 2.17 x 10^-13^, **Supplemental Figure 7**). We repeated this for mouse liver and heart TF ChIP-seq peaks [39] and obtained similar results for the default FIMO motif hit cutoff (liver p = 2.81 x 10^-78^, heart p = 7.65 x 10^-65^), a stricter FIMO motif hit cutoff (liver p = 3.03 x 10^-25^, heart p = 5.22 x 10^-65^), and the relationship between motif hit q-values and peak signals (liver p = 1.10 x 10^-3^, heart p = 2.90 x 10^-4^), suggesting that the relationship between the presence of the downstream motif and CTCF binding strength is consistent across tissues.

## DISCUSSION

We developed a new approach for discovering *in vivo* TF binding motifs for different DBDs within a TF by training and interpreting a neural network for predicting whether a wild-type TF ChIP-seq peak will be significantly stronger than the corresponding peak from mutated TF ChIP-seq data. We applied this approach to a dataset with ChIP-seq from wild-type and mutated CTCF and, in addition to identifying the known motifs of CTCF ZFs 3-11, we discovered a putative interaction between CTCF ZF 1 and a novel downstream GAGCCA motif that as well as a putative weak interaction between CTCF ZF 2 and an ATT motif connecting the core and discovered downstream motifs (**Figure 4**). Our discovered downstream motif is supported by *in vitro* studies of CTCF from HT-SELEX [31] and EMSA [36] and *in vitro* ZF studies from B1H assays [33, 35], the weak putative motif for ZF 2 was supported by CTCF HT-SELEX data, and the discovered downstream motif occurs more frequently in CTCF ChIP-seq peaks that are not bound by CTCF-s than by those that are. While, to fully demonstrate the validity of our findings, we would need to experimentally test if ZFs 1-2 can interact with these sequences using an assay like EMSA, our combination of existing and novel approaches for using additional datasets to support our findings provide a foundation for computationally following up on potential motif discoveries.

**Figure 4:**
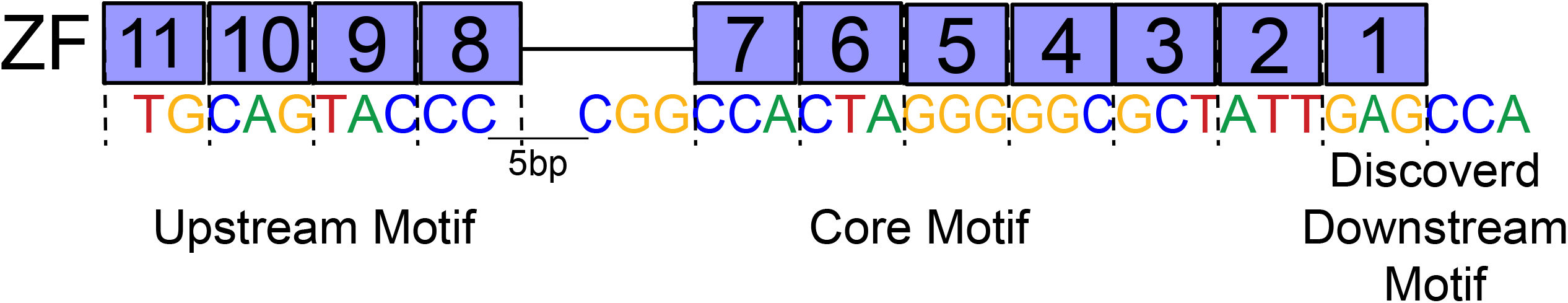
Proposed Motif for CTCF Based on Findings from Interpreting Neural Networks.

Our modeling approach enabled us to discover both known and novel motifs as well as the spacings between them because a neural network interpreted with deepLIFT [28] followed by TF-MoDISco [29] does not require an explicit featurization of the sequence or assumptions about the sizes of the motifs and because the tasks for our neural networks directly contrasted the wild-type and mutant datasets. Some previous studies have used k-mer support vector machines (SVMs) [40, 41], but linear SVMs cannot identify relationships between nucleotides that span more than k bases, and k needs to be small (usually at most eleven) so that the number of parameters does not become too large to be learned with the available data, making these models incapable of learning our longer motifs (**Figure 2b**). Many additional studies have trained neural networks to predict TF binding and used interpretation methods similar to those that we used to discover known and sometimes novel motifs of TFs [42–47], but the subset of these studies that predicted CTCF binding failed to identify our motif for ZFs 1-2, likely because, unlike our study, their models were not designed to directly learn individual DBD binding preferences. In fact, a previous study suggested that, for TFs with multiple ZFs, some ZFs have consistent binding patterns across the majority of binding sites, while others bind at only a minority of sites and do not always have the same spacing when binding, making their motifs difficult to detect when modeling all TF binding sites together [48]. Properly evaluating differences between wild-type and mutant TF binding requires multiple high-quality replicates of *in vivo* binding data from each of a wild-type and mutant TF, which, unfortunately, are not always available; this inability to properly detect differential binding due to generating only one biological replicate may explain why the only existing studying contrasting wild-type and mutant CTCF binding [49] was unable to obtain some of our results (**Supplemental Notes**).

Our modeling approach has several drawbacks beyond requiring *in vivo* binding data from a mutated TF. One limitation is that our negative set consisted of a combination of the peaks that were comparably strong in the wild-type and the mutant and peaks that were stronger in the mutant, preventing us from identifying motifs that are associated with destabilizing interactions between ZFs in CTCF and DNA. We think this is why we did not identify the downstream motif identified by the previous study of this dataset [11], a motif that is thought to destabilize the binding of ZFs 2-3 to DNA. One possible way to extend our neural network to handle this case would be to train and interpret a regression model for predicting the fold-change of the wild-type versus the mutant peak strengths. In addition, convolutional neural networks have inherent limitations, regardless of how their tasks are defined, that may limit our ability to use our interpretations of them to decipher DBD binding (**Supplemental Notes**). Our ability to discover potentially novel motifs for CTCF’s ZFs in spite of the limitations of our approach demonstrates that our approach provides a foundation for identifying motifs of TF DBDs.

## CONCLUSIONS

To our knowledge, we are the first to train machine learning models to predict whether a wild-type TF will have stronger binding than a mutated TF and the first to use differential binding between a wild-type and mutated TF to decipher binding preferences of the TF’s DBDs. Our approach can aid future comparisons of wild-type TF binding to binding of TFs whose DBDs have been mutated, including TFs whose motifs are not well-characterized. In addition, our approach could be extended to comparisons of other *in vivo* TF binding experiments, such as differential TF binding across conditions, cell types, or time points.

## METHODS

### CTCF ChIP-seq Data Processing

We reprocessed the ChIP-seq data from wild-type CTCF and each of the CTCF ZF mutants so that we could ensure that it met ENCODE quality control standards after applying recently recommended methods for filtering reads and identifying reproducible peaks [50, 51]. To do this, we downloaded the data from GSE33819 [11, 52, 53]. We then mapped reads to mm10 [54] and filtered reads using the AQUAS Transcription Factor ChIP-seq processing pipeline [55] with default parameters.

To ensure that our datasets were sufficiently high-quality for reliable downstream analysis, we used the AQUAS pipeline [55] with default parameters to perform strict quality control evaluations. We first evaluated whether a dataset had more signal than we would expect from reads randomly dispersed in the genome, which we did by computing the normalized strand coefficient (NSC), which should ideally be at least 1.05, and the relative strand correlation (RSC), which should ideally be at least 0.8 [50]. We found that all of the datasets had NSC > 1.05 and RSC > 0.8. Since all of the biological replicates for each mutant met ENCODE standards [50], we did not remove any datasets for our analyses.

Before calling peaks, we combined the reads into “pooled replicates,” which are all of the reads pooled across the three biological replicates. For the wild-type dataset, we used the tagged data from *Mus musculus* so that the species and experimental protocol would be consistent with those of the mutants; a previous study showed that the peaks from the tagged data are consistent with those from a CTCF antibody ChIP-seq experiment done in the same lab [11]. Then, for each dataset, we used the AQUAS pipeline [55] to randomly divide the reads from each dataset into two “pooled pseudo-replicates,” which are groups containing half of the reads, so that we could later evaluate the reproducibility of our peaks. We then used the AQUAS pipeline [55] with default parameters to call peaks and identify irreproducible discovery rate (IDR) reproducible peaks [56] across pooled pseudo-replicates. We identified reproducible peaks across pooled pseudo-replicates because we had three biological replicates for each dataset, and we wanted to incorporate data from all of them. We obtained tens of thousands of IDR reproducible peaks for the wild-type and for each mutant.

### Identifying Differential Peaks

To identify differential peaks, which we defined as peaks that are significantly stronger in the wild-type than they are in the mutant, we merged peaks from the different datasets, computed the number of reads from each dataset in each merged peak, and evaluated whether the read depth was significantly larger in the wild-type than in each mutant. We merged all IDR reproducible peaks from each mutant, including the R339W mutant for ZF 3, and the tagged *Mus musculus* wild-type by merging peaks whose summits were within 50bp of each other and defining the merged peak summit to be the average of the summits of the combined peaks. Next, we used pybedtools version 7.10.0 [57, 58] and to remove reads from each replicate of each experiment mapping to mitochondrial DNA, unknown chromosome, or random chromosome parts; shift reads to the right by half of their fragment lengths from cross-correlation analysis; and count the reads overlapping the five-prime end of each merged peak. Finally, we ran DESeq2 [27] on the read counts to compare peaks in the wild-type to those in each mutant. We defined a peak to a member of the positive set, meaning significantly stronger in the wild-type, if the q-value was less than 0.05 and the log base 2 fold-change was greater than 1 and a member of the negative set if the log base 2 fold-change was less than or equal to 0.

### Training Neural Networks for Differential Peak Prediction

For each mutant except for R339W, which was not thought to have a substantial effect on binding [11], we trained a separate neural network to predict whether a merged peak was a member of the positive or negative set. Merged peaks that were members of neither set were not used. For each merged peak, we created two examples: the sequence underlying the merged peak summit +/- 500bp and the sequence underlying the reverse complement of the merged peak summit +/- 500bp. Our training set was chromosomes 3-7, 10-19, and X; our validation set was chromosomes 8-9; and our test set was chromosomes 1-2. We one-hot-encoded the sequences as four-by-one thousand matrices, where each row contained a binary vector indicating whether each position in the sequence consisted of a specific nucleotide; this encoding method has been used in previous studies that applied neural networks to predict TF binding [59–61]. We encoded N’s as all zero’s. Thus, our input data did not contain any prior information about what parts of the DNA sequence are involved in CTCF binding.

The architecture that we used for our neural network was three convolutional layers [25], which were each followed by a rectified linear unit, followed by a max-pooling layer. The convolutional filters on the first layer should identify motifs that reveal whether a peak is significantly stronger in the wild-type, the filters in the following layers should identify combinations of those motifs, and the max-pooling layer encodes the assumption that a single motif combination should not occur multiple times within a short region. The first convolutional layer had 60 4 x 15 filters with stride 1 x 1, the second convolutional layer had 60 1 x 15 filters with stride 1 x 1, and the third convolutional layer had 15 1 x 15 filters with stride 1 x 1. Each layer had dropout rate 0.2. The max-pooling layer was size 1 x 35 with stride 1 x 35. The max-pooling layer was followed by fully connected layer with a sigmoid output. The neural networks were trained using Keras version 0.3.2 [62] and Theano version 0.8.2 backend [63] using stochastic gradient descent with Nesterov momentum 0.85 [64] and learning rate 0.01, batch size 200, and class weights set to the fraction of peaks in the other class. These hyperparameters were selected after evaluating performance of multiple sets of hyperparameters on the validation set. The early-stopping criterium was three consecutive epochs with no improvement in recall at eighty percent precision on the validation set. Weights were initialized to be those from a pre-trained neural network with the same hyper-parameters and the negative set randomly down-sampled to be the size of the positive set. The weights for the pre-training were initialized using Keras’s He normal initializer [62, 65].

### Identifying Important Features Learned by Neural Networks for Differential Peak Prediction

Motifs that are important for making correct positive predictions are likely to be indicative of the binding preference of the mutant ZF because they are important for determining whether a peak will be significantly stronger in the wild-type data than in the data from TF in which that ZF was mutated. To identify these motifs, we computed the importance of every nucleotide in each true positive example in the validation set and then used these importance values to construct motifs. We scored the importance of every nucleotide in every true positive example in the validation set using deepLIFT, which computes the contribution of each nucleotide to a sequence’s prediction relative to a reference [28]. We used the deepLIFT version 0.5.5-theano with the Rescale rule, where scores were taken from the sequence layer with the target of the final convolutional layer and our reference was a sequence of N’s. We used an extension to deepLIFT with the Rescale rule to compute the “hypothetical scores,” which can be thought of the extent to which the classifier is expecting a nucleotide, for each nucleotide at each position in each sequence [29].

The deepLIFT scores and hypothetical scores were inputted into the TF-MoDISco method for constructing “TF-MoDISco motifs” learned by the model [29]. TF-MoDISco first extracts sequence patterns that frequently have high deepLIFT scores in ChIP-seq peak sequences (called seqlets), next computes the pairwise similarities between seqlets, and then uses the similarities cluster the seqlets into “TF-MoDISco motifs.” We ran TF-MoDIsco with these settings: seqlet FDR threshold = 0.2; gapped k-mer settings for similarity computation k-mer length = 8, number of gaps = 1, and number of mismatches = 0; final motif width = 50; and minimum number of seqlets = 200. We used the aggregated hypothetical scores of the seqlets supporting each TF-MoDISco motif to construct motif images.

To make position frequency matrices from TF-MoDISco motifs, we averaged the one-hot-encoded sequences at all of the seqlet coordinates associated with the motifs. We also extracted the upstream, core, and discovered downstream motifs from our TF-MoDISco motifs (**File S1**). To extract the upstream motif, we removed degenerate positions from the ends of the TF-MoDISco motif for ZF 11. We did this by first identifying the upstream-most position in which at least one nucleotide had probability > 0.60 and removing all earlier positions. We then scanned the motif until reaching another position at which no nucleotides had probability > 0.60. Because the following position was non-degenerate, we continued searching for an additional position in which no nucleotides had probability > 0.60. We removed that and all further downstream positions in the TF-MoDISco motif. To extract the core motif, we repeated the same process that we used for the upstream motif on the TF-MoDISco motif from ZF 6, except that we used a probability cutoff of 0.40 and required two consecutive bases with nucleotides passing the probability cutoff to begin extracting the motif. To extract the discovered downstream motif, we repeated the process that we used for the upstream motif on the TF-MoDISco motif from ZF 1, except that we used a probability cutoff of 0.35 and started at the downstream end of the TF-MoDISco motif, scanning upstream towards the start; we stopped when the difference in nucleotide probability for the nucleotide with the greatest probability decreased by > 0.35 between two consecutive positions. We used these upstream, core, and downstream motifs for further analyses. Finally, we constructed six motifs, which we call “mega-motifs”: the core motif (**Supplemental File 1**); the upstream motif (**S2 File**); the discovered downstream motif (**S3 File**); the upstream motif followed by the core motif, where the motifs were separated by seven bases with nucleotide probabilities corresponding to the G/C-content in mouse (The upstream and core motifs we identified were separated by seven bases because the nucleotide probabilities of two most upstream bases of the known core motif were not large enough to be captured in our core motif.); the core motif followed by the discovered downstream motif, where the motifs were separated by two bases with nucleotide probabilities corresponding to the G/C-content in mouse (The core and discovered downstream motifs we identified were separated by two bases.); and the upstream motif followed by the core motif followed by the discovered downstream motif, where the upstream and core motifs were separated by seven bases with nucleotide probabilities corresponding to the G/C-content in mouse and the core and discovered downstream motifs were separated by two bases with nucleotide probabilities corresponding to the G/C-content in mouse.

### Logistic Regression with Motif Hit Scores

We compared the performance of our neural network to that of a logistic regression with the scores of motif hits of the three motifs from [11]. We received the three motifs described in [11] in MEME format [66] from the authors of [11]. We scanned the merged CTCF peaks for these motifs using FIMO version 4.12.0 [38] with default parameters, where the background was the background provided to us by the authors of [11]. We computed the smallest motif q-value in each peak for each motif and used the negative log base ten of that q-value as a feature in a logistic regression; if there were no motif hits with q-value < 0.5 for a motif in a peak, then we set the value of that feature to zero for that peak. We trained the logistic regression using Scikit-learn version 0.19.1 [67] with l2 penalty 1.0. We used the same positives and negatives that we used for our neural network. We trained the logistic regression on a combination of the training and validation sets that we used for our neural network and evaluated the logistic regression using the same test set that we used for our neural network. Note that the original motifs and spacings between them were found using all of the peaks in from the wild-type, including those on the chromosomes that we held out for testing; thus, we may be underestimating the difference in performance between our neural networks and the logistic regressions with the original motif hit scores.

We also compared the performance of our neural network and of the logistic regression with the original motif hit scores to the performance of a logistic regression where the only feature was the top TF-MoDISco motif (TF-MoDISco motif with the most supporting seqlets) score and to a logistic regression in which the only feature was the score of the original upstream motif followed by five base pairs followed by the original core motif. For the latter, the nucleotide frequencies in the five base pairs between the original upstream and original core motifs were set to be the background single nucleotide frequencies provided by the authors of [11]. For both of these evaluations, we computed features and trained logistic regressions using the same procedures that we used for the logistic regressions with the original motif hit scores.

### Area Under Precision-Recall Curve Computation

We compared the performances of the logistic regressions with motif hit scores to those of our neural networks by computing the area under the precision-recall curve for each model. We computed this using PRROC [68]. We used this metric instead of AUROC because our negative set is always larger than our positive set (**Supplemental Table 1**).

### Identifying Motif Combinations in Reads from CTCF HT-SELEX Data

We compared the core motif followed by the discovered downstream motif to reads from CTCF HT-SELEX data from [31]. We first downloaded the reads from cycle 0 (control), which were taken before the TF was introduced, and cycle 4, the final cycle, that were generated for CTCF HT-SELEX in [31]. Since the HT-SELEX reads were only 20bp long, we constructed a partial combination of the core motif followed by the discovered downstream motif, which was the downstream 10bp of the core motif followed by 2bp with the G/C-content in mouse (the core and discovered downstream motif were separated by 2bp) followed by the upstream 4bp of the downstream motif. We then scored the motif match to each HT-SELEX read by converting the read and its reverse complement into a one-hot-encoded sequence, computing the dot product of those sequences and the partial combination of the core motif followed by the discovered downstream motif at every possible alignment of the two matrices, and computing the maximum of the dot products. We compared the distribution of scores for reads from cycle 0 to the distribution of scores for reads from cycle 4 using a Wilcoxon rank-sum test; the histograms of these distributions are illustrated in **Figure 3b**. As a control, we repeated this process only the downstream 16bp of the core motif, and the histograms for this comparison are in **Supplemental Figure 4**.

We created aggregate motifs by running FIMO [38] with the partial combination of the core motif and the discovered downstream motif on reads from CTCF HT-SELEX cycle 4 [31], one-hot-encoding the positions with motif hits, averaging the one-hot-encoded matrices, and visualizing these averages as motif logos (**Supplemental Figure 5**). We defined a “motif hit” to motif hits with FIMO q-value less than four different cutoffs – 0.05, 0.01, 0.005, and 0.001 – and created an aggregate motif for the motif hits from each of these cutoffs. We visualized the motif logos using meme2images from the MEME suite [66].

### Comparison of CTCF Peaks Overlapping CTCF-s Peaks to Those that Do not Overlap CTCF-s Peaks

To compare the CTCF peaks that overlap CTCF-s peaks to those that do not, we re-processed that biotin-tagged data from [24], identified motif hits of the core motif followed by the discovered downstream motif in the CTCF ChIP-seq peaks, and compared the p-values of the motif hits in different subsets of the peaks. We re-processed the data and evaluated data quality using the AQUAS pipeline [55] with the hg38 genome assembly [69] and default parameters; both the CTCF and CTCF-s data had NSC > 1.05 and RSC > 0.8. Unless otherwise indicated, we used IDR reproducible peaks across pseudo-replicates (Each dataset had only 1 biological replicate.) for our analyses. We identified motif hits of the core motif followed by the discovered downstream motif in the CTCF ChIP-seq peaks by first using bedtools [58] to obtain the fasta file for the peaks, next running the MEME suite’s fasta-get-markov [66] with -m 1 on the fasta file to obtain a background file, and then running FIMO [38] on the fasta file with the background file and the core followed downstream mega-motif (**Supplemental File 1**) with settings --max-stored-scores 50000000 and --thresh 1. We used bedtools intersect with settings -wa and -u to obtain CTCF ChIP-seq peaks that overlap CTCF-s ChIP-seq peaks, and we used bedtools subtract with setting -A to obtain CTCF ChIP-seq peaks that do not overlap any, including non-reproducible across pseudo-replicates, CTCF-s ChIP-seq peaks. We then obtained the p-value of the best core followed by downstream mega-motif match in each of these subsets of CTCF ChIP-seq peaks, setting the p-value to 1 when no motif match was identified. We compared the distributions of the p-values for these two subsets of CTCF ChIP-seq peaks using a Wilcoxon rank-sum test.

To investigate whether our results could be explained by a relationship between core followed by discovered downstream motif occurrences and reproducibility of CTCF ChIP-seq peaks across experiments, we also compared CTCF ChIP-seq peaks from [24] to those from ENCODE [32]. Since the data in [24] came from HeLa cells, we downloaded the “optimal” IDR reproducible peaks (ENCODE entry ENCFF772LNY) and peaks from pooled reads across replicates (ENCODE entry ENCFF331BAX) from the deepest ENCODE HeLa cell CTCF ChIP-seq dataset [32]. We used bedtools intersect with settings -wa and -u to obtain CTCF ChIP-seq peaks from [24] that overlap ENCODE IDR reproducible CTCF ChIP-seq peaks, and we used bedtools subtract with setting -A to obtain CTCF ChIP-seq peaks from [24] that do not overlap ENCODE pooled replicate CTCF ChIP-seq peaks. We then obtained the p-value of the best core followed by downstream mega-motif match in each of these subsets of CTCF ChIP-seq peaks from [24], setting the p-value to 1 when no motif match was identified. We compared the distributions of the p-values for these two subsets of CTCF ChIP-seq peaks from [24] using a Wilcoxon rank-sum test.

### Computational Predictions of CTCF Motifs Using Models Trained on *in vitro* ZF Binding Data

To further evaluate whether our discovered downstream motif is likely to interact with CTCF ZF

1, we compared it to predicted CTCF motifs from models trained using *in vitro* ZF binding data. These models were trained on *in vitro* data measuring the binding specificities of individual ZFs; they take ZF amino acid sequences as input and output a predicted motif. First, we used RCADE2’s RC.sh to predict the motif for CTCF [35]. To explore alternative methods, we also put the sequences of CTCF’s ZFs into the “Predict PWMs” function of the “Interactive PWM Predictor” [33, 34]. We predicted the motif using each of the three available models: “RF Regression on B1H,” “Expanded Linear SVM,” and “Polynomial SVM.” We additionally ran each model on ZF 1 alone to confirm that the models predicted that ZF 1 interacts with GAG. **Figure 3a** contains the outputs from RCADE2, and **Supplemental Figure 6** contains the outputs from the other models.

### Comparison of CTCF ChIP-seq Peak Strengths with Different Combinations of Motifs

We compared peak strengths for different motif combinations by identifying occurrences of each mega-motif in CTCF peaks, grouping peaks based on mega-motif presences, and quantifying properties of each peak in each group. We scanned the wild-type mouse CTCF peaks for the mega-motifs using FIMO [38] with default parameters except for the threshold, which we set to 1, and the background, which we set to the output from fasta-get-markov [66] run on the sequences of the CTCF peaks with setting -m 1. We used version 4.12.0 of the MEME suite [66] for all of these analyses.

We evaluated the relationship between peak strength and the presence of the downstream motif by comparing peaks with the core followed by downstream mega-motif to peaks with only the core motif. We defined motif hits as motif occurrences with FIMO p-value < 0.0001 (default from FIMO) [38]. When using the stricter motif cutoff, we defined motif hits as motif occurrences with FIMO q-value < 0.05. We then used bedtools version 2.26.0 [58] to identify peaks with the core motif, the core motif and no discovered downstream motif, and the core followed by downstream mega-motif for the different motif hit cutoffs. We defined the peak strength to be the natural log of the signal from SPP (column seven from the narrowPeak files). We compared the peak strength for pair of peaks – peaks with the core motif and no discovered downstream motif versus peaks with the core followed by downstream mega-motif – by doing a two-sided Wilcoxon rank-sum test, and we did a Bonferroni correction of the p-values by multiplying them by six (two comparisons for each of three cell types/tissues). We repeated this process for liver and heart data, which was taken from the mouse ENCODE 8-week-old mouse Ren Lab datasets [39].

Since the definition of a motif hit can be sensitive to thresholding, we also compared the peak strength of CTCF peaks to the −log base ten q-values from FIMO [38] of all of the motif occurrences from FIMO regardless of their FIMO p-value or q-value. We incorporated all motif occurrences by identifying the correlation between peak strength and the −log base ten q-values from the FIMO for the core motif and the core followed by downstream mega-motif, and the upstream followed by core mega-motif. We then compared the correlations for the core motif and each of the other two mega-motifs using a one-sided Fisher’s r-to-z transformation and did a Bonferroni correction of all p-values by multiplying them by six (two pairs for each of three tissues).

## DECLARATIONS

### Availability of data and materials

All data used in this study was previously published and released in other studies. Mouse activated B Cell CTCF ChIP-seq data analyzed in this study was downloaded from GEO GSE33819 [11]. CTCF HT-SELEX data was downloaded from ENA PRJEB3289 [31]. Mouse liver and heart CTCF ChIP-seq data were downloaded from the ENCODE portal entries ENCFF542WEE and ENCFF616HYA, respectively [39]. Corresponding CTCF and CTCF-s ChIP-seq data were downloaded from GSE108869. ENCODE HeLa cell CTCF ChIP-seq data was downloaded from ENCODE portal entry ENCSR000AOA [32]. The zinc finger image in **Supplemental Figure 1** was taken from [70]. The core motif logo in **Figure 1** and **Supplemental Figure 1** is the Hocomoco human CTCF motif downloaded from CIS-BP [17], and the upstream motif in **Supplemental Figure 1** is from [11]. Code can be found at https://github.com/kundajelab/CTCFMutants, and additional code can be made available from the authors upon request. Core, upstream, and discovered downstream motifs are in **Supplemental File 1**. Deep neural network models, deepLIFT scores, TF-MoDISco motifs, and FIMO hits for motifs can be found on the **Supporting Website**.

## Acknowledgements

We would like to thank A. Kundaje, A. Shrikumar, H. B. Fraser, the other members of the Fraser and Kundaje Labs, R. Casellas, J. Tycko, and Z. Zuo for useful discussions and suggestions. We would also like to thank the members of the Casellas Lab for providing us with the original motifs from [11] and the background used for tools in the MEME suite from that study.

## Authors’ information

Study conceptualization was done by I.M.K. and C.S.F. Data curation was done by I.M.K. Resource management and methodology development were done by I.M.K. with assistance from C.S.F. Formal analysis, investigation, software implementation, and visualization of computational results were done by I.M.K. with assistance from A.B. Original draft preparation was done by I.M.K. Reviewing and editing was done by all authors.

