## Supplemental Information for "Neural network modeling of differential binding between wild-type and mutant CTCF reveals putative binding preferences for zinc fingers 1-2"

**CONTENTS**


1. Supplemental Notes
2. Supplemental Figure Captions
3. Supplemental Tables
4. Additional Supplemental Information
5. Supplemental References
6. Supplemental Figures

**SUPPLEMENTAL NOTES**

**Comparison of Our Study to Recent Study Comparing Wild-Type and Mutated CTCF Binding**

A recent study did ChIP-seq on wild-type CTCF in mouse embryonic stem cells as well as CTCF with deletions of ZFs 1, 8, 9, 10, and 11 and, in addition to finding that deleting ZF 8 led to weakening of topologically associated domains and changes in DNA methylation and gene expression, found a weak motif for ZF 1 that has some similarity to our discovered downstream motif [1]. While this study did investigate peaks that are substantially weaker in the wild-type than in the mutant, the study had only one replicate for each mutant and for the wild-type, limiting its ability to reliably detect differential binding. In contrast, the dataset that we used had multiple replicates for the wild-type and for each mutant, allowing us to identify differential binding events that are unlikely to be explained by differences in when an experiment was done [2]. In addition, the previous study identified motifs associated with ZFs by aggregating the sequences in the differential regions, while we identified specific nucleotides that are predictive of CTCF binding being significantly stronger in the wild-type. The previous study not only failed to confidently identify our motif for ZF 1 but found that the motif ZF 8 is almost identical to the motifs for ZFs 9-11, while our motif for ZF 8 placed less importance than our motifs for ZFs 9-11 on the most upstream part of the upstream motif, matching the known biology that different ZFs interact with different parts of their TFs' motifs [3].

**Limitations of Neural Networks**

Neural Networks also have inherent limitations, regardless of how their tasks are defined. For example, neural networks require a large number of training examples [4], so they may not always be usable for mutants that do not affect at least a few thousand peaks, and they may not be able to learn motifs that are not present in at least a few thousand peaks in the training set. This may explain why our neural network was unable to identify the slight decrease in the frequency of C relative to the T in the eighth position of the core motif identified by a previous study of the peaks that are lost in a ZF 1 mutant [5]. In addition, machine learning models may learn the minimal set of features that are necessary for achieving good predictive performance; as a result, if there are multiple highly correlated features that are associated with the model's task, the model may learn only a strict subset of them, so the motifs learned by the model may exclude some biologically relevant motifs. In addition, convolutional neural networks require a fixed-size input [6], which is why we used merged peak summits +/- 500bp. Using a model that can handle inputs of variable sizes would enable us to incorporate additional information that has been proposed to affect TF binding, such as sequences of distal regions that loop to TF peaks. Recent advances have enabled sequences of variable sizes to be used as inputs to deep convolutional neural networks [7, 8], so such modeling may be achievable. Finally, the failure of the logistic regression with the TF-MoDISco motif hit score to reach the performance of our neural network and the lack of additional meaningful TF-MoDISco motifs for the neural networks for the mutants of ZFs 1-8 suggest that our methods for interpreting what our neural networks learned are suboptimal; thus, improving neural network interpretation methods should enable us to use neural networks to discover additional novel biology.

#### SUPPLEMENTAL FIGURE CAPTIONS

**Supplemental Figure 1: What We Expected the Neural Networks (NNs) to Learn Based on Previous Studies**

We obtained zinc finger images from [9]. The core motif logo in this figure is the Hocomoco human CTCF motif downloaded from CIS-BP [10], and the upstream motif is from [11].

**Supplemental Figure 2: Test Set Area Under the Precision-Recall Curve (AUPRC) of Motif Hit Score Logistic Regressions for the Original Upstream Motif Followed by the Original Core Motif versus Neural Networks and Top TF-MoDISco Motif Hit Score Logistic Regressions**

**Supplemental Figure 3: Top Two TF-MoDISco Motifs for Mutations in Zinc Figures 9-11**

The top two TF-MoDISco motifs for **a)** mutation in ZF 9, **b)** mutation in ZF 10, and **c)** mutation in ZF 11 are the upstream followed by the core motif with two different spacings, where the top-ranked TF-MoDISco motif (most supporting seqlets) has the more common spacing according to previous studies, and the second highest-ranked TF-MoDISco motif (second most supporting seqlets) has the less common spacing according to previous studies. The tick marks indicate the nucleotide positions. The core motif logo in this figure is the Hocomoco human CTCF motif downloaded from CIS-BP [10], and the upstream motif is from [11].

**Supplemental Figure 4: Comparison of Motif Hit Scores of the Core Motif in Reads from CTCF HT-SELEX Data in Cycle 0 to Cycle 4.**

**Supplemental Figure 5: Comparison of TF-MoDISco Motifs from the Mutants of ZFs 1 and 2 to Aggregated Reads from CTCF HT-SELEX Cycle 4 with Matches at Different q-Value Cutoffs**

We truncated TF-MoDISco motifs to the 16bp that align to the parts of the core and downstream motifs, which we used to identify motif matches in the HT-SELEX reads.

**Supplemental Figure 6: Comparison of the TF-MoDISco Motif from the Mutant of ZF 1 to Computationally Predicted Motifs of CTCF's DBDs**

We compared the TF-MoDISco motif from the mutant of ZF to computationally predicted motifs of CTCF's DBDs from three different models – "Interactive PWM Predictor RF Regression on B1H," "Interactive PWM Predictor RF Expanded Linear SVM," and "Interactive PWM Predictor RF Polynomial SVM," – trained on *in vitro* B1H ZF binding data [12, 13].

**Supplemental Figure 7: Comparison of Ctcf Peak Strengths with Motif Hit Scores for Different Motif Combinations**

Correlations between wild-type Ctcf ChIP-seq peak strength and negative log base ten of the motif hit q-values from FIMO (illustrated as density plots). Correlations are the Pearson correlation, and p-value is from the Fisher's r-to-z test with a Bonferroni correction.

#### SUPPLEMENTAL TABLE

**Supplemental Table 1: Number of Positives and Negatives in the Training Set for Each Model**

| Mutant Zinc Finger | Number of Positives in Training Set | Number of Negatives in Training Set |
|---|---|---|
| 1 | 19,916 | 152,810 |
| 2 | 19,708 | 161,390 |
| 3 | 67,620 | 151,486 |
| 4 | 68,906 | 142,768 |
| 5 | 69,054 | 146,680 |
| 6 | 84,120 | 136,944 |
| 7 | 80,778 | 141,906 |
| 8 | 22,312 | 147,102 |
| 9 | 52,358 | 148,766 |
| 10 | 35,134 | 156,456 |
| 11 | 41,360 | 146,400 |

#### ADDITONAL SUPPLEMENTAL INFORMATION

**Supplemental File 1: Motifs Extracted from deepLIFT Scores Using TF-MoDISco**

**Supplemental Website:** http://mitra.stanford.edu/kundaje/imk1/CTCFMutantsProject/

1. **Results from DESeq2 and corresponding peak summits:**
   http://mitra.stanford.edu/kundaje/imk1/CTCFMutantsProject/DESeq2Results
2. **Deep neural network weights and architectures:**
   http://mitra.stanford.edu/kundaje/imk1/CTCFMutantsProject/DeepNeuralNetworkModels
3. **hdf5 and bigwig files with deepLIFT scores and maximum deepLIFT scores at each nucleotide for each neural network:**
   http://mitra.stanford.edu/kundaje/imk1/CTCFMutantsProject/DeepLIFTScores
4. **TF-MoDISco results and full set of TF-MoDISco motifs for all neural networks:**
   http://mitra.stanford.edu/kundaje/imk1/CTCFMutantsProject/TFMoDIScoMotifs
5. **Results from FIMO on wild-type peaks:**
   http://mitra.stanford.edu/kundaje/imk1/CTCFMutantsProject/WT_rep1-pr.IDR0.05.filt.FIMOResultsNewTFModiscoMotifsAllHits

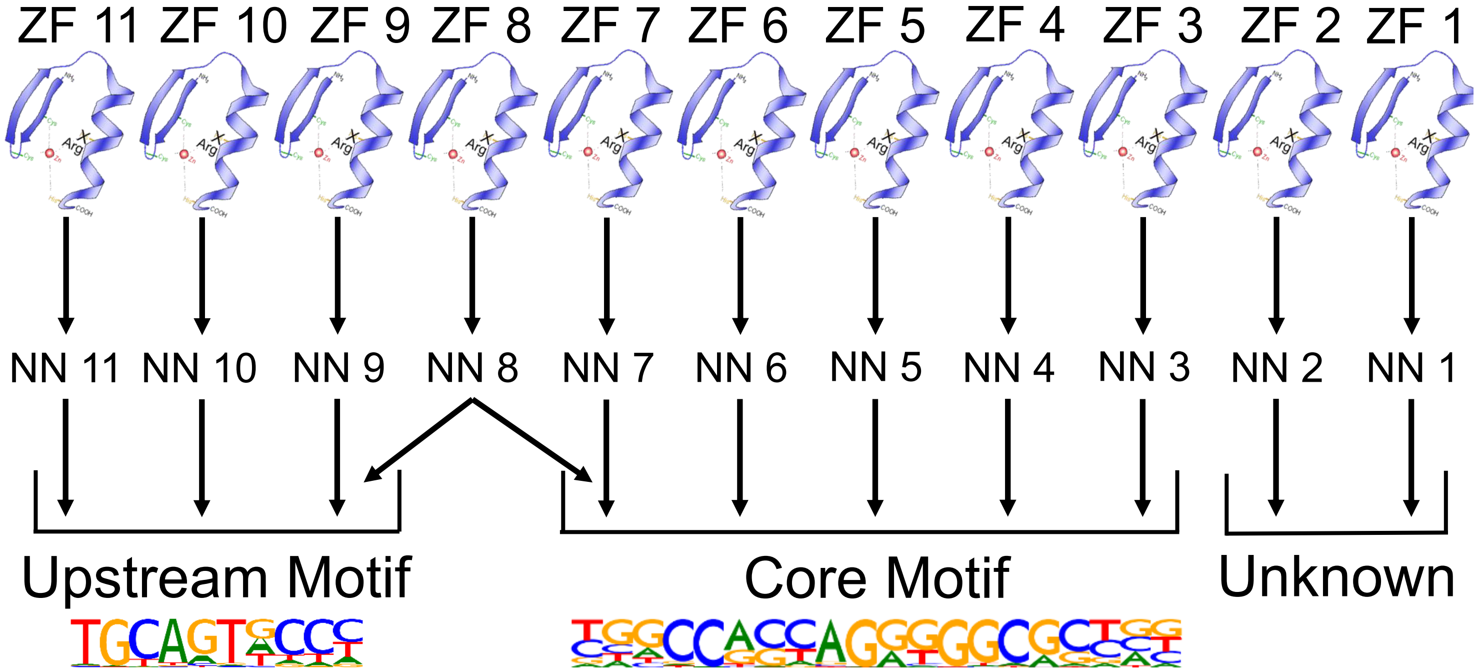


**Supplemental Figure 1**

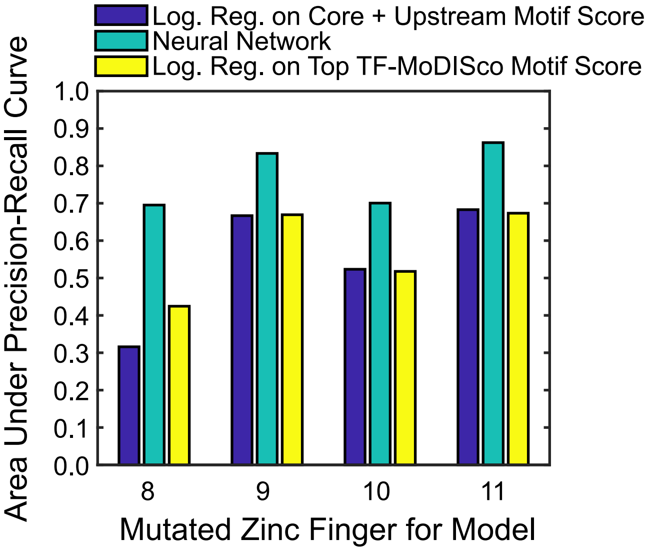


### Supplemental Figure 2

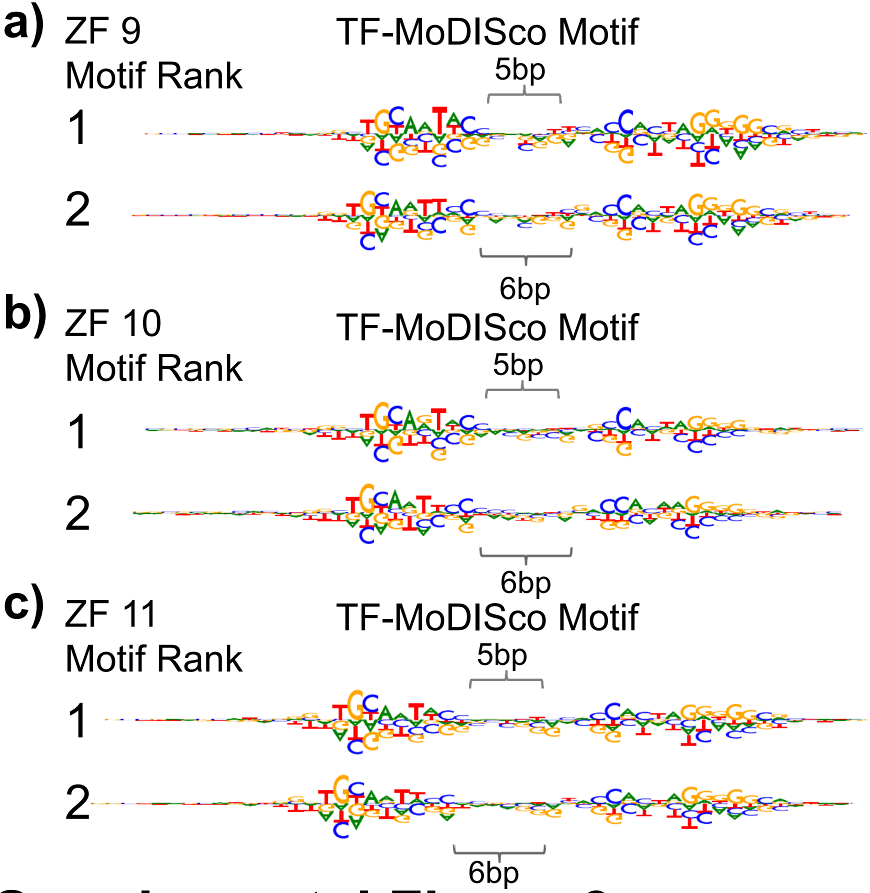


### Supplemental Figure 3

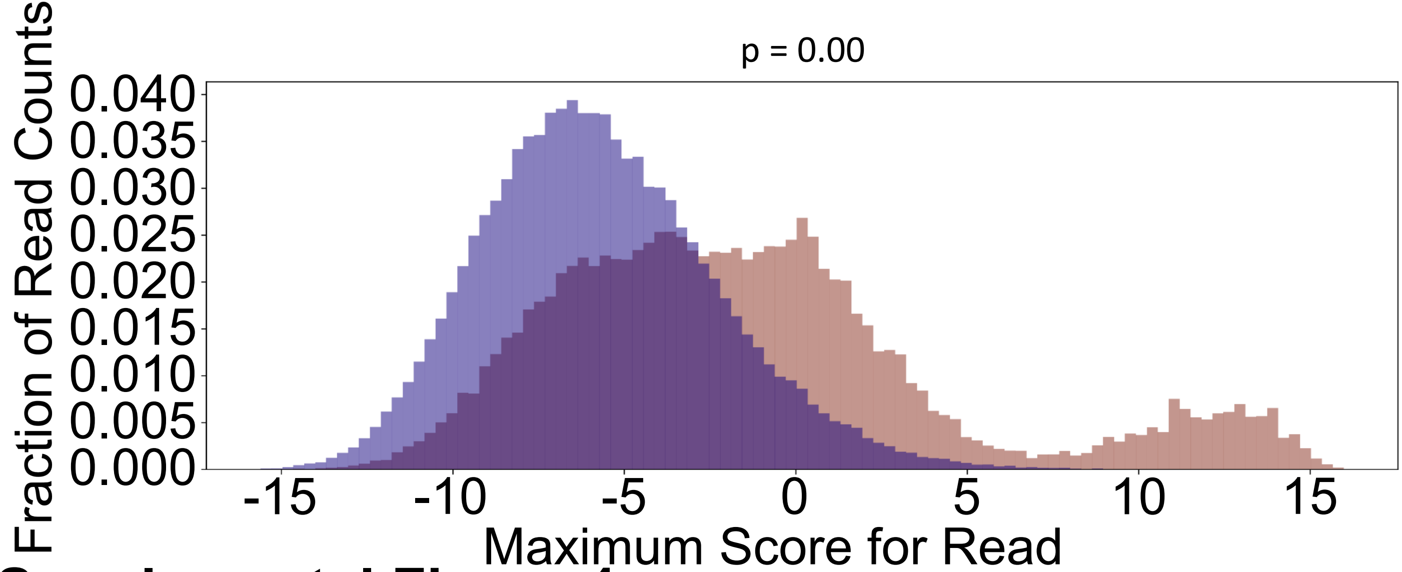


**Supplemental Figure 4**

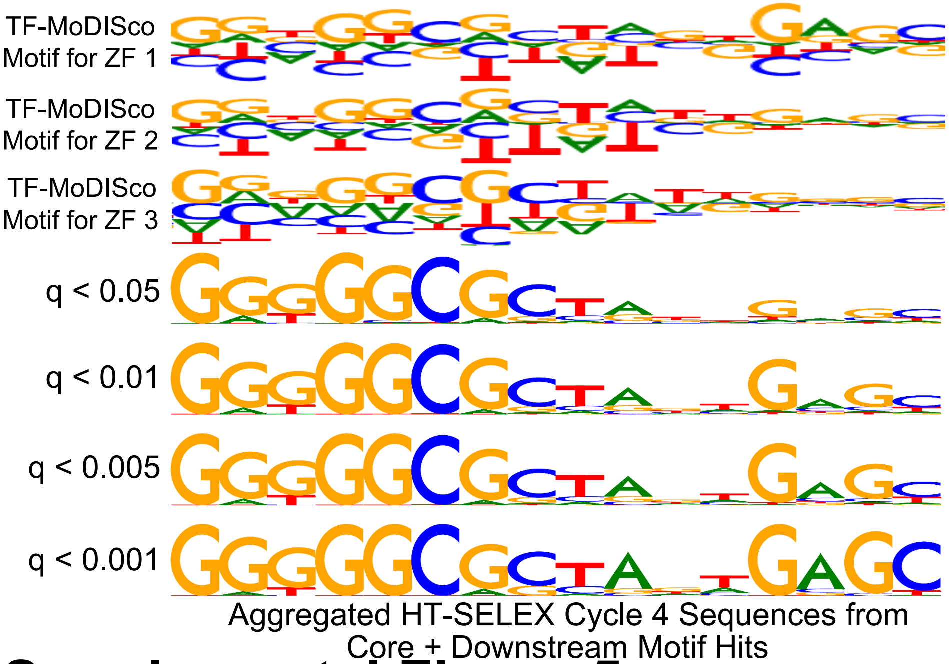


### Supplemental Figure 5

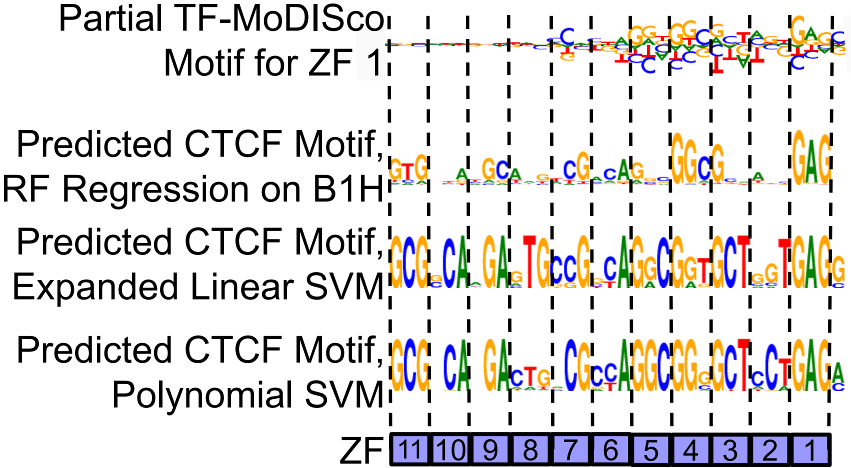


### Supplemental Figure 6

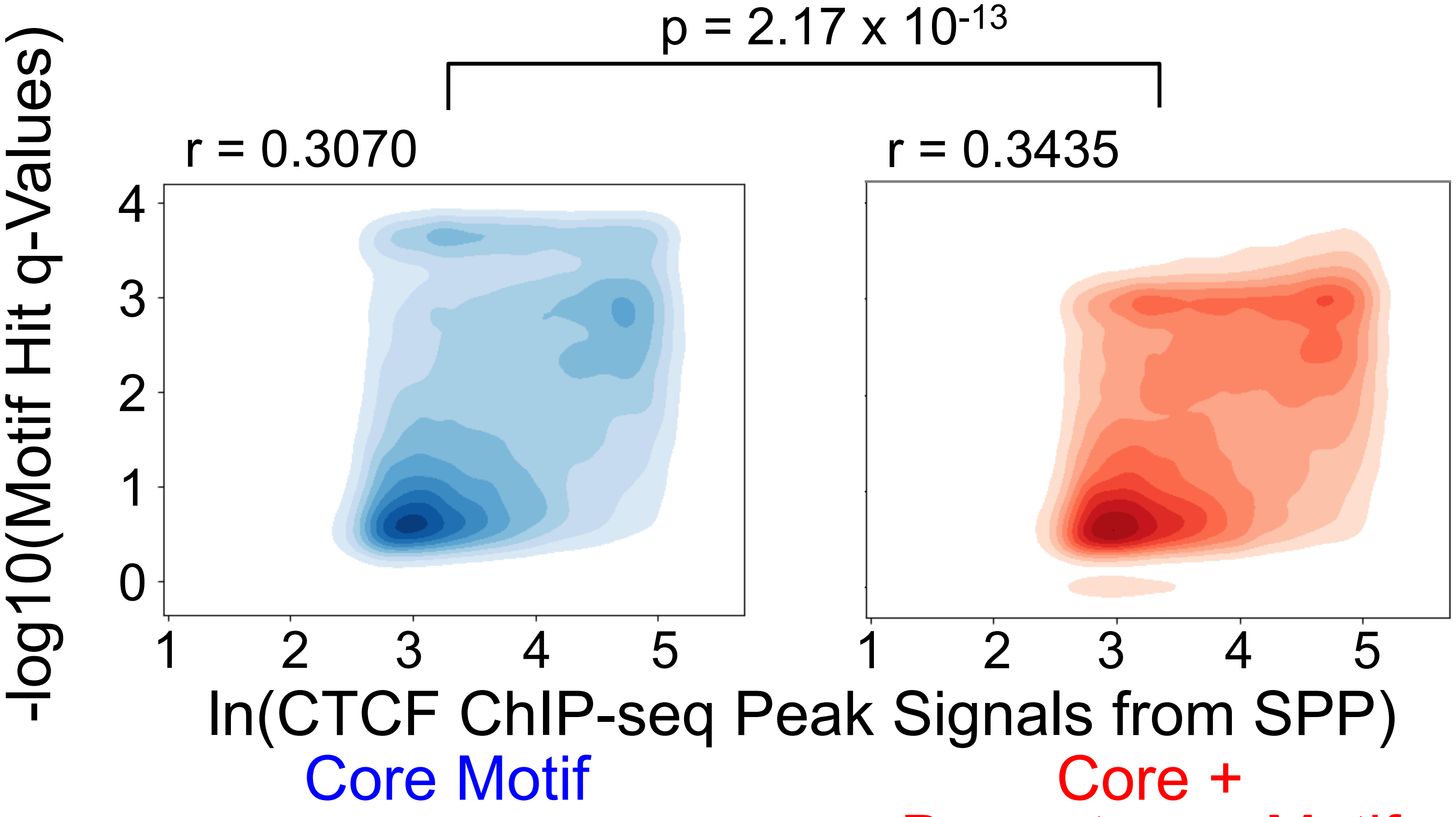


**Supplemental Figure 7**
